# Heterogenization of remaining biodiversity in fragmented tropical forests across agricultural landscapes

**DOI:** 10.1101/629782

**Authors:** Cristina Yuri Vidal, Diogo Souza Bezerra Rocha, Marinez Ferreira de Siqueira, Ricardo Ribeiro Rodrigues, Tadeu Siqueira

## Abstract

The increasing worldwide interest on the conservation of tropical forests reflects the conversion of over 50% of their area into agricultural lands and other uses. Understanding the distribution of remaining biodiversity across agricultural landscapes is an essential task to guide future conservation strategies. To understand the long-term effects of fragmentation on biodiversity, we investigated whether forest fragments in southeastern Brazil are under a taxonomic homogenization or heterogenization process. We estimated pre-deforestation species richness and composition based on a Species Distribution Modelling approach, and compared them to the observed patterns of α- and β-diversity. In particular, we asked (i) if changes in β-diversity reveal convergence or divergence on species composition; (ii) if these changes are similar between forest fragments in Strictly Protected Areas (SPAs) (n=20) and within private lands (n=367) and in different regions of the state (West, Center, and Southeast). We detected steep reductions in observed local species richness in relation to our modeled predictions, and this was particularly true among forest fragments in non-protected private lands. The higher observed β diversity indicated an overall biotic heterogenization process, consistent with the idea that the originally diverse vegetation is now reduced to small and isolated patches, with unique disturbance histories and impoverished communities derived from a large regional species pool. Since conservation of biodiversity extends beyond the boundaries of strictly Protected Areas, we advocate forest fragments are valuable for conservation in agricultural landscapes, with particular relevance for private lands, which represent the most exposed and neglected share of what is left.

## INTRODUCTION

Tropical forests are known for holding a substantial portion of the world’s terrestrial biodiversity (Myers et al. 2000; Chazdon, Harvey, et al. 2009; Slik et al. 2015), yet over 50% of its original area was converted into agricultural lands or other uses, with prospects for an increasing agricultural expansion in tropical developing countries (Hansen et al. 2013; Laurance et al. 2014). Understanding the distribution of remaining biodiversity across human-modified landscapes (HMLs) and what has changed after forest conversion are essential questions to guide future conservation strategies (Tabarelli et al. 2010; Melo, Arroyo-Rodríguez, et al. 2013; Laurance et al. 2014; Socolar et al. 2016). In HMLs, tropical forest fragments comprehend a variety of different sized habitats, including forests that never experienced clear cutting or severe impacts (*i.e.* primary forests), and the full spectrum of degraded forests that are regenerating after extraction, fire or abandonment of croplands and pastures, among other previous land-uses (*i.e.* secondary forests) (Melo, Arroyo-Rodríguez, et al. 2013; Malhi et al. 2014). A very narrow fraction of these forest fragments are under restrictive protected areas, where biodiversity conservation is tangible to a limited extent (Andam et al. 2008; Joppa et al. 2008; Coetzee et al. 2014). In this context, the variety of forest fragments located within private lands not only represent the largest share of what is left (Gardner et al. 2009; Sparovek et al. 2012; Soares-Filho et al. 2014; Mendenhall et al. 2016), but also the most neglected. These fragments are rarely explicitly targeted in conservation programs, as the focus is usually on avoiding deforestation (Chazdon, Harvey, et al. 2009; Barlow et al. 2016). Several studies have shown that secondary forests may play an important role in conservation (Santos et al. 2007, Chazdon et al. 2009; Dent & Wright 2009; Tabarelli et al. 2012), as they hold a depleted but relevant portion of biodiversity even within HMLs. Abundant evidence is available for birds (Karp et al. 2012; Emer et al. 2018), mammals (Galetti et al. 2009; Pardini et al. 2010; Beca et al. 2017) and plants (Arroyo-Rodríguez et al. 2008; Lima et al. 2015; Sfair et al. 2016; Farah et al. 2017).

The efects of habitat loss and fragmentation on biodiversity have been intensively studied over the past three decades, with a primary focus on more local scales (Karp et al. 2012; Vellend et al. 2013; Dornelas et al. 2014; Malhi et al. 2014; Newbold et al. 2015; Barlow et al. 2016). There has been an increase on studies focusing on broader extensions, based on the assumption that we cannot properly understand the consequences of deforestation if disregarding the influence of entire landscapes over local processes (Malhi et al. 2014). Also, beyond the intuitive interest on species richness loss, an emerging issue of interest is how community composition responds to fragmentation along spatial gradients and periods of time (Arroyo-Rodríguez et al. 2013; Solar et al. 2015; Morante-Filho, Arroyo-Rodríguez, et al. 2015; Collins et al. 2017; Olden et al. 2018). Measures of the variation in species composition among sites (β) can indicate if communities are converging or diverging in response to fragmentation, providing relevant information on the maintenance of regional diversity (Socolar et al. 2016).

Some studies have demonstrated that forest fragmentation and degradation result into biotic homogenization (Vellend et al. 2007; Lôbo et al. 2011; Karp et al. 2012; Marcelo Tabarelli et al. 2012; Püttker et al. 2015; Zwiener et al. 2017), *i.e.* the convergence of biotas in time and space, in which communities may suffer a simplification of their genetic, taxonomic and functional diversities (McKinney & Lockwood 1999; Olden & Rooney 2006). The rationale is that more ecologically specialized species (“losers”) are locally extinct, while a much narrower sub-set of generalists, with high dispersal abilities (“winners”), override them (Lôbo et al. 2011; Marcelo Tabarelli et al. 2012; Mendenhall et al. 2016). This process results in impoverished communities that represent sub-sets of a larger pool of species, translated by reduced β-diversity and high contribution of the nestedness component on β diversity. The predominance of nestedness suggests conservation efforts might focus on the richest sites, provided they are connected to support viable communities (Howe 2014; Socolar et al. 2016; Emer et al. 2018). An opposite consequence to fragmentation and degradation occurs when communities diverge on composition over time and space (*i.e.* enhanced β diversity) because they suffer different frequencies and levels of disturbances combined with dispersal limitations and environmental heterogeneity, resulting into biotic heterogenization (Dornelas et al. 2014; Solar et al. 2015; Sfair et al. 2016; Catano et al. 2017; Collins et al. 2017). In this scenario, widespread regional diversity conservation is only possible if targeting multiple sites.

Very few studies of plant community changes in response to fragmentation evaluate β-diversity patterns based on temporal replicates (Lôbo et al. 2011; Dornelas et al. 2014; Haddad et al. 2015; Collins et al. 2017); most of them adopt a space-for-time approach (*e.g.*, disturbed × undisturbed) regardless of the fact that distinct sites could reflect distinct pre-disturbance conditions (Collins et al. 2017). In addition to the lack of temporal replicates, severely deforested landscapes may not be suitable for the space-for-time approach when in the absence of large forest remnants or high forest-covered regions to represent undisturbed ecosystems. For that matter, environmental niche modeling (ENM) and species distribution modeling (SDM) can be useful to provide species’ spatial occurrence disregarding the effects of anthropogenic disturbances, which is a major driver of community composition changes (Malhi et al. 2014, Catano et al. 2017). Since ENM is based on the niche concept and considers environmental conditions as the primarily influence over the establishment of a given species (De Marco Junior & Siqueira 2009), it results in maps representing the geographic space where the abiotic conditions are appropriate (Peterson & Soberon 2012). Distinctly from ENM, which disregards dispersal/colonization limitations and biotic interactions, SDM restrict the model calibration to accessible areas, incorporating dispersal issues into analyses and producing maps where a focal species may potentially occur, with varying degrees of suitability (De Marco Junior & Siqueira 2009; Peterson & Soberon 2012). For community-level modeling, further methodologies are available to adjust the over-prediction of the number of species coexisting at a given location (Guisan & Rahbek 2011; Calabrese et al. 2014; Gavish et al. 2017). Therefore, these approaches combined can be used to generate expectations of what communities would look like, in terms of species composition, if they were not disturbed by land use conversion.

In recent years, Brazil has stood out among tropical developing countries for its environmental engagement, which resulted on the exceptional decline in deforestation rates during 2000-2012 (Hansen et al. 2013; Loyola 2014), despite the fact that it already has 30% of its total area occupied by agricultural lands (Martinelli et al. 2010). In Sao Paulo state, southeastern Brazil, which includes two biodiversity hotspots (Atlantic Forest and Cerrado) (Myers et al. 2000; Laurance 2009), deforestation has taken place during the last three centuries (Metzger 2009), and surveys only became a common practice in the last 30 years (Haddad et al. 2015; Renato Augusto Ferreira de Lima et al. 2015). With a very long history of land conversion for agricultural purposes, most remaining vegetation is comprised by small forest fragments (i.e. <50ha) (Ribeiro et al. 2009), representing an unique opportunity to understand the long-term effects of fragmentation on biodiversity. The purpose of our study was to evaluate whether the woody assemblages on forest fragments are under a taxonomic homogenization or heterogenization process in response to habitat fragmentation. For that matter, we estimated pre-deforestation species richness and composition based on a Species Distribution Modelling approach, and compared them to the observed patterns of α- and β-diversity. In particular, we asked (i) if changes in β-diversity indicate convergent or divergent composition; (ii) if these changes are similar between forest fragments under strict protection or within private lands and in different regions of the state. In the hyper-fragmented landscapes of this study, we expected to find lower mean values of α-diversity within private lands relative to strictly Protected Areas. Additionally, because of intrinsic environmental heterogeneity strengthened by fragmentation disturbances, we expected an overall increase in β-diversity, indicating a taxonomic differentiation process. This pattern should be particularly more evident for unprotected forest fragments located within private lands, where fragments are more susceptible to a broader range of recurrent disturbances (Laurance et al. 2014; Malhi et al. 2014).

## METHODS

### Study region

The state of Sao Paulo is located within the range of two current global hotspots, the Atlantic Forest and Cerrado (tropical savannas) (Myers et al. 2000). With a long history of deforestation caused by timber extraction and agricultural cycles (coffee, pasture, orange, sugarcane) (Metzger 2009), the São Paulo state case-study may provide relevant insight about the long-term effects of habitat loss and fragmentation on biodiversity, which can be useful for other tropical regions facing the same threats (Laurance 2009). Both hotspots are poorly protected, with only 1.6% and 0.5% of Atlantic Forest and Cerrado’s original area protected as strictly Protected Areas (Durigan et al. 2006; Ribeiro et al. 2009; Carranza et al. 2014). The remaining vegetation cover in the interior plateau (*i.e.* excluding coastal areas) ranges from 1 to 30% (São Paulo State Forest Inventory 2011), mostly located within private rural properties (Gardner et al. 2009; Sparovek et al. 2012; Soares-Filho et al. 2014; Mendenhall et al. 2016).

In order to facilitate analyses’ interpretation along the extent region of this study, we adopted the ecological regions defined by Setzer (1966), which divide the state in 6 sub-regions based on climate, soil, topography and vegetation variables (Setzer 1966). We excluded the south and north coastal areas because their forest cover is well above the rest of the state. To meet a minimum of 30 localities per sub-region, we joined Southwest and Northwest into one single “West” sub-region (**Figure 1**).

**Figure 1:**
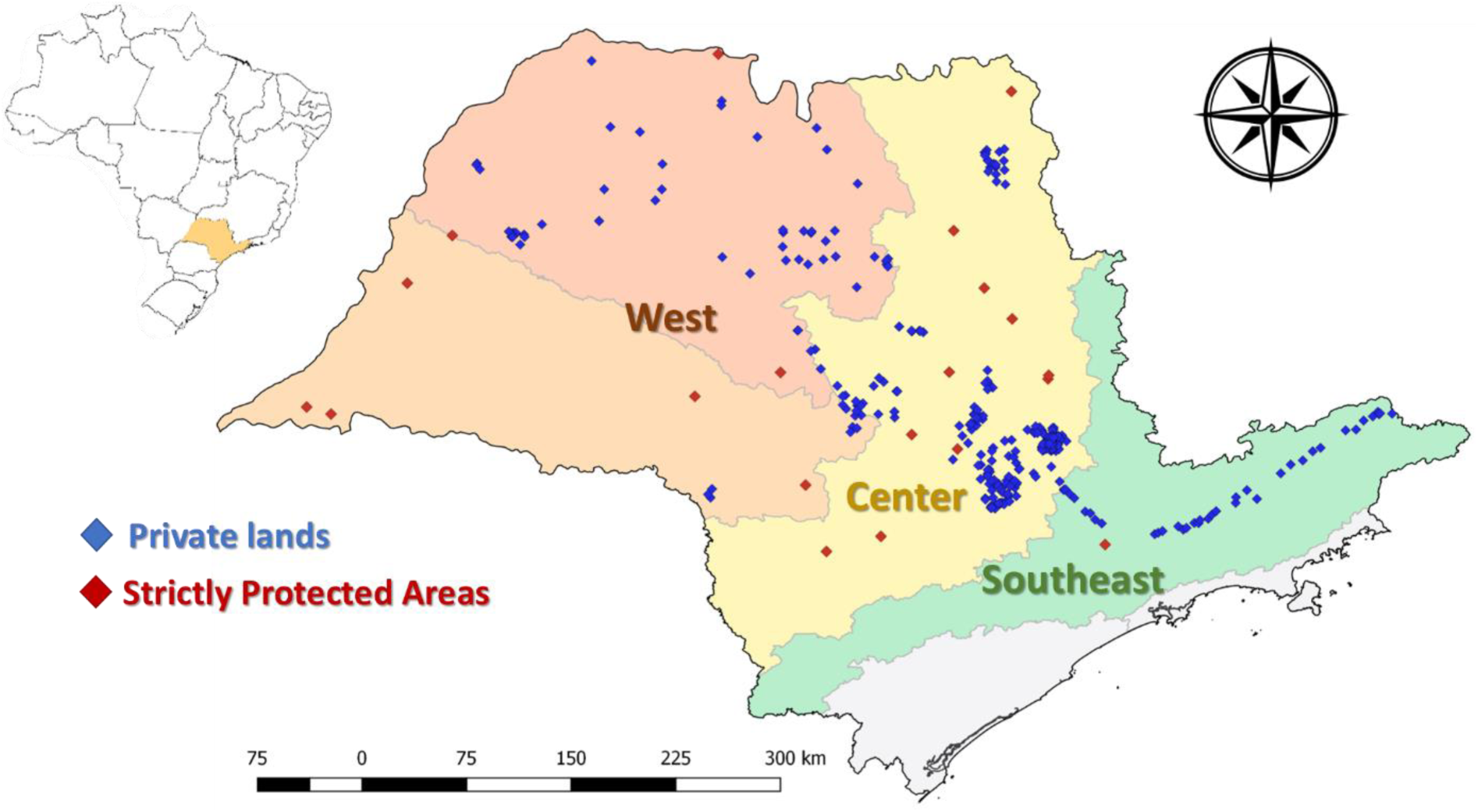
Distribution of forest fragments among regions in São Paulo state (Southeastern Brazil), considering those in private lands (n=367) and strictly protected areas (n=20). Mean forest cover based on São Paulo State Forest Inventory (2011): West = 6.5%, Center= 10.9%, Southeast= 27.7%.

Regarding vegetation, we focused on the predominant seasonal semi-deciduous forest (SSF), considering its transition to evergreen forests or forested savannas (Cerradão) (Oliveira-Filho & Fontes 2000; Durigan & Ratter 2006), and all other forest ecosystems included in this extension: swamp, alluvial and deciduous forests. Despite the fact that these forests are influenced and determined by soil, altitude and climatic conditions (Oliveira-Filho & Fontes 2000), their floristic composition are strongly influenced by the surrounding vegetation (*e.g.* SSF), creating complex transitional mosaics and continuum distribution of species (Kurtz et al. 2015; Oliveira-Filho & Fontes 2000).

### Woody plant species occurrence data

We chose to use woody plant species (*i.e.* trees and shrubs) in this study because they represent a fundamental structure and functional component of forest ecosystems, as they support food webs and represent a substantial proportion of tropical diversity (Arroyo-Rodriguez et al. 2015; Slik et al. 2015). Based on a compilation of 367 floristic surveys in private lands (more details in Rodrigues et al. 2011 and Farah et al. 2017), we initially defined a species pool of 921 species. To enhance our sample of the environmental space occupied by these species and to improve modelling outcomes, we retrieved complementary national occurrence data for these species from *SpeciesLink* (http://splink.cria.org.br/) and NeoTropTree (http://prof.icb.ufmg.br/treeatlan/) (Oliveira-Filho 2017). Finally, we selected 691 species occurring in more than 10 localities along the Brazilian territory, which comprised 114,740 records.

We also gathered available checklists for 20 strictly Protected Areas within the study region (**Figure 1**) following the same criteria regarding forest types as those applied to select forests in private lands. We did not use the strictly Protected Areas localities to build species distribution models due to the lack of precision on their geographical coordinates, varying from a random point within their boundaries to the municipality centroid. We compiled the floristic surveys available at Fundação Florestal (http://fflorestal.sp.gov.br) and Instituto Florestal (http://iflorestal.sp.gov.br), both related to the São Paulo State Environmental Secretariat, and WWF Protected Areas’ Observatory (http://observatorio.wwf.org.br/). Species names were standardized using the Plantminer web tool (www.plantminer.com) (Carvalho et al. 2010), based on Flora do Brasil (www.floradobrasil.jbrj.gov.br) (Flora_do_Brasil_2020) and The Plant List (www.theplantlist.org/). Complementary queries were performed on The Missouri Botanical Gardens (www.tropicos.org). According to these databases, we excluded any exotic and unidentified species from final compilation.

### Environmental data

We compiled 22 environmental predictors with spatial resolution of 1km^2^ and summarized them by using a Principal Component Analysis (PCA) considering 1,000 randomly distributed points within Sao Paulo state. Pairs of variables with scores > I1I (absolute value) were verified to avoid multicollinearity (correlation <0.7) finally selecting 6 variables with the highest PCA scores (**Appendix 1**).

### Environmental Niche Modeling and Species Distribution Modeling

Environmental Niche Models (ENM) are statistical models that relate focal species occurrence to associated environmental conditions, generating correlative rules that allow extrapolation and prediction of occupancy patterns over wide geographic extents, representing a valuable tool for conservationist purposes (De Marco Junior & Siqueira 2009; Angelieri et al. 2016; Gavish et al. 2017; Guisan et al. 2013). We applied a Species Distribution Model (SDM) approach by restricting ENM to accessible areas, aiming to presume the distribution range of species if they were not affected by habitat loss, fragmentation and disturbance. In other words, if species’ distribution were primarily defined by abiotic conditions (*i.e.*, environmental niche), in the lack of constraints imposed by altered habitat and landscape structure (*e.g.*, fragmentation and patch isolation) (Peterson & Soberon 2012). As ENM and SDM can be based on the same sets of mathematical algorithms, occurrence data and environmental variables (Peterson & Soberon 2012), they are near-synonymous: the main difference is that SDM implies on some sort of restriction over ENM, which will be further detailed; for the purpose of this study, we hereafter will refer to our modeling approach only as SDM.

We built the SDM for each species using the Model-R framework (Sánchez-Tapia et al. 2018) with a three-fold cross validation procedure, meaning that two partitions were used for parameter estimation and algorithm training, and one to evaluate the model’s accuracy. Random pseudo-absence points (n_back_ = 1000) were sorted within a mean distance buffer, where the radius of the buffer was the mean geographic distance between the occurrence points. If one species’ records were less than 20 km apart, they were rarefied to reduce effects of sampling bias and avoid modelling overfitting (Elith et al. 2006; Zwiener et al. 2017).

In the Model-R framework, for each partition and algorithm a model was built and its performance was tested by their True Skill Statistics (TSS) (Allouche et al. 2006). We previously tested several algorithms – BioClim, GLM, SVM, Random Forest, MaxEnt – and selected the last two based on their overall performance (**Appendix 2**), which was consonant to the results found by Diniz-filho et al. (2009). We then selected Random Forest and MaxEnt partitions with TSS>0.4, and applied a threshold that maximizes two error types: sensitivity (*i.e.* true presences) and specifity (*i.e.* true absences) (Sánchez-Tapia et al. 2018). The resulting binary models were averaged into a final model for each algorithm, an then combined into a final ensemble model with an average threshold that maximizes TSS values (Sánchez-Tapia et al. 2018), resulting in a final map indicating areas of probable presence. These maps were primarily generated for all the Brazilian territory and then cropped for São Paulo State.

### Species richness and community composition

Our analyses were based on two different type of information: the observed species richness and community composition in each site and the species richness and community composition predicted by SDM. The observed metrics were adjusted to consider only modelled species, that is, species occurring in more than ten localities, at least 20km apart and with a final ensemble model derived from Random Forest and Maxent algorithms, using their partitions with TSS>0.4 (**Figure 2a**), as previously detailed. This adjustment was necessary in order to make proper comparisons between observed and predicted richness and compositions, and considered a final sub-set of 663 woody species.

**Figure 2:**
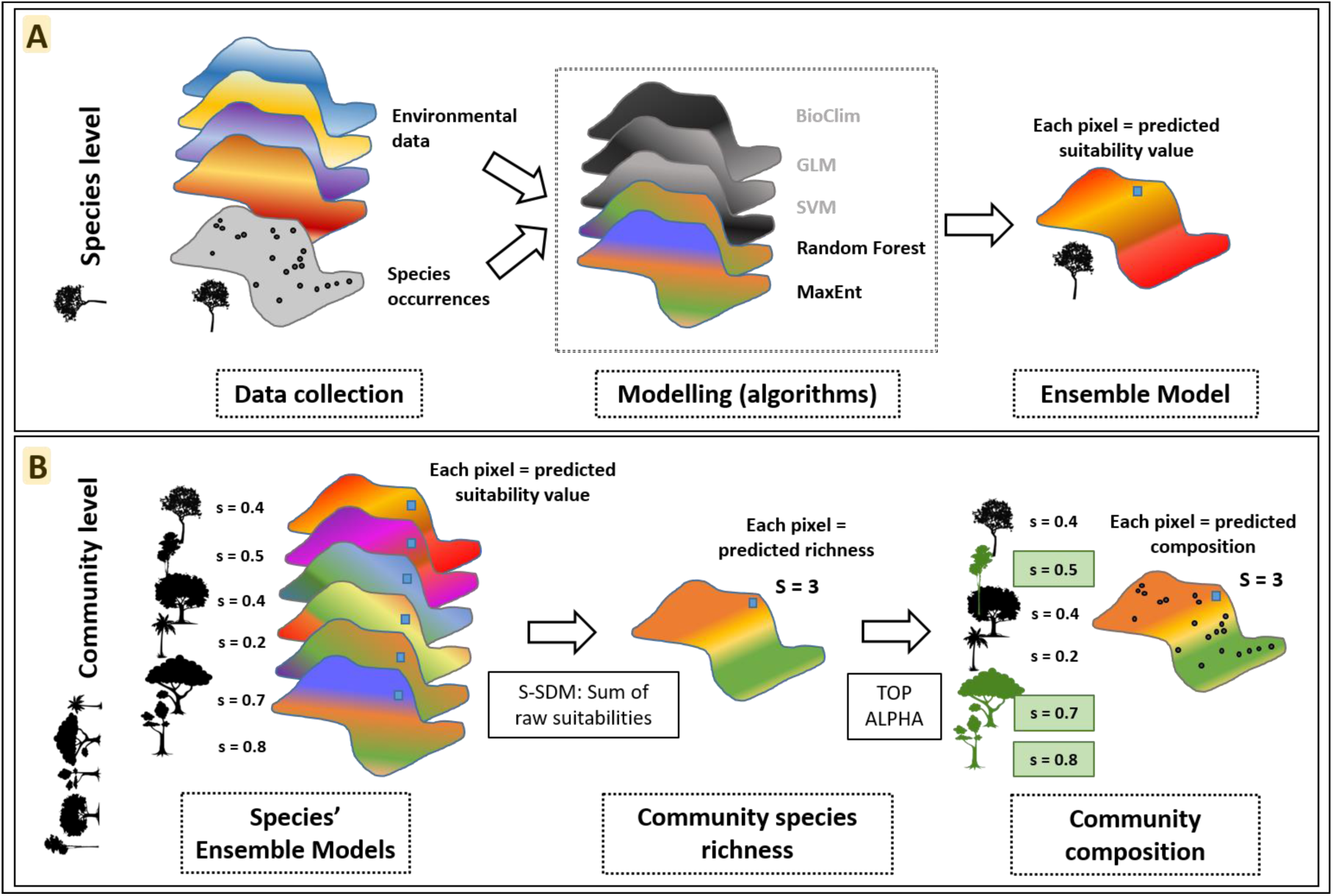
Steps for species distribution modeling and for predicting community richness and composition at a given point (i.e. pixel). (A) Step 1: Species distribution modeling for each of the 663 species, considering Random Forest and MaxEnt algorithms and their partitions with True Skill Statistics (TSS) ≥0.4. The final ensemble model indicate, for each pixel, the suitability of occurrence for one given species. (B) Step 2: The predicted community species richness at a given point results from the sum of the raw probabilities of occurrence (*i.e.*, suitabilities not converted to binary values by any threshold) of targeted species. Step 3: The community composition results from the TOP ALPHA approach, which consists on ranking the species according to their suitabilities of occurrences and selecting species with the highest values until attaining the predicted richness (step 2).

Species richness based on SDM can be predicted either by stacking individual species-level models (Stacking SDM, S-SDM) or by modelling α-diversity itself (Macroecological Models, MEM) (Gavish et al. 2017; Calabrese et al. 2014; D’Amen et al. 2015; Guisan & Rahbek 2011). Since stacking binary presence/absence SDM tend to overpredict richness, as it does not account for biotic interactions or filters (Gavish et al. 2017; Guisan & Rahbek 2011), Calabrese *et al.* (2014) proposed S-SDM corrections to reduce these overpredictions and concluded that if stacked correctly, S-SDM are no worse than MEM. According to their findings, a corrected S-SDM approach involves summing-up the raw predicted suitabilities for each locality instead of summing-up their binary values (Calabrese et al. 2014; D’Amen et al. 2015) (**Figure 2b**). Hence, we inputted an average of the raw suitability values into the areas of probable presence defined on the final ensemble models; the predicted community species richness is the sum of these values considering any given location (i.e. pixel).

Following the estimation of potential richness, we predicted site-level composition by adopting the “top alpha” approach (Gavish et al. 2017), where we ranked the species’ suitabilities of occurrence per site from the highest to the lowest values – based on their individual ensemble SDM – and then selected the top number of species that equals to the predicted potential richness per site (D’Amen et al. 2015; Gavish et al. 2017) (**Figure 2b**).

### β-diversity analyses

The compositional variation among communities from site-to-site (β-diversity) relates local diversity (α) to the regional species pool (γ) (Anderson et al. 2010). When evaluating the effects of habitat loss and fragmentation, changes on the organization of biodiversity over space and time can reveal if biological homogenization or heterogenization is taking place, an essential information to guide conservation planning over regional diversity (Socolar et al. 2016; Arroyo-Rodríguez et al. 2013; Püttker et al. 2015). Despite the valuable contribution from evaluating β-diversity, its interpretation must be very cautious, as there are several ways to measure and compare it (Koleff et al. 2003; Baselga et al. 2007; Jost 2007; Chao et al. 2012; Jost et al. 2010; Anderson et al. 2006, 2010; Tuomisto 2010).

There is extensive debate regarding the interrelationships among α, β and γ-diversity, in addition to measures for partitioning it (*i.e.* multiplicative or additive) and statistical approaches to properly analyze β-diversity (Anderson et al. 2010). A particular concern for our study is the fact that a great variety of metrics to estimate β-diversity depend on α and γ diversity – and therefore on scale and sample size. Considering that our samples represent a compilation of floristic surveys using distinct methods and sampling effort, we decided to use a β-diversity metric that weights on composition dissimilarities more than on richness differences (Koleff et al. 2003). For that matter, we calculated pairwise Sorensen (**β**_**SOR**_) and Simpson indices (**β**_**SIM**_) among sites, which indicate the overall variation on the species composition between pairs of sites (**β**_**SOR**_) and the variation related to its turnover component (**β**_**SIM**_), reflecting the replacement of species (Baselga 2010; Baselga et al. 2015; Socolar et al. 2016).

To evaluate differences between observed and predicted β-diversity, we: (i) ran a Principal Coordinate Analysis (PCoA) based on the Simpson (**β**_**SIM**_) observed dissimilarity matrix (as the turnover component was the main contribution to total beta diversity; see bellow); (ii) estimated the distance of each site (forest fragment) to the group centroid in the multivariate ordination space generated by the PCoA (**β**_**d**_, **Figure 3**); (iii) repeated steps (i) and (ii) using the SDM predicted species composition; and (iv) compared mean observed **β**_**d**_ with mean predicted **β**_**d**_ with permutational paired t-test. In this test, observed **β**_**d**_ values were paired with predicted **β**_**d**_ ones. **β**_**d**_ is analogous to the local contribution to beta diversity (LCBD) proposed by Legendre and De Cáceres (2013), since higher values of **β**_**d**_ represent higher distinctiveness of one site (forest fragment) within a group. This metric with standard effect size allowed us to test the null hypothesis that beta diversity does not differ among observed and predicted communities.

**Figure 3:**
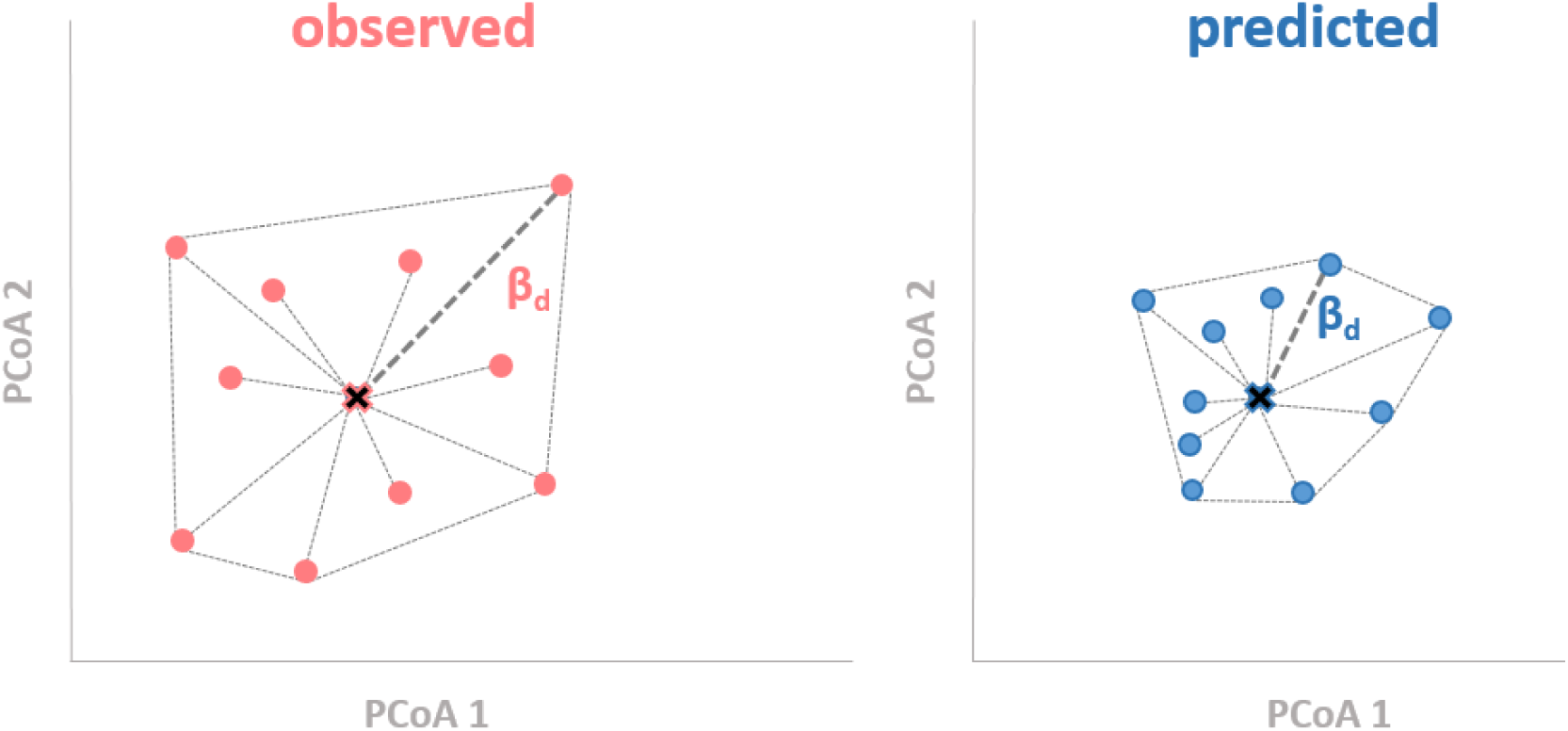
Hypothetical example of a multivariate ordination space describing the distance (**β**_**d**_) of a unit to the group centroid (central cross), considering the observed (red) and the predicted (blue) species composition.

## RESULTS

For the 663 woody species considered in this study, overall comparisons revealed that predicted richness at the site level (α-diversity) was 3.8 times higher than observed richness. This ratio was much lower when considering only strictly Protected Areas, ranging around 1.0. In fact, the few forest fragments that presented increased observed species richness in relation to predicted richness were mostly represented by Protected Areas (**Figure 4**) (**Appendices 3 and 4**).

**Figure 4:**
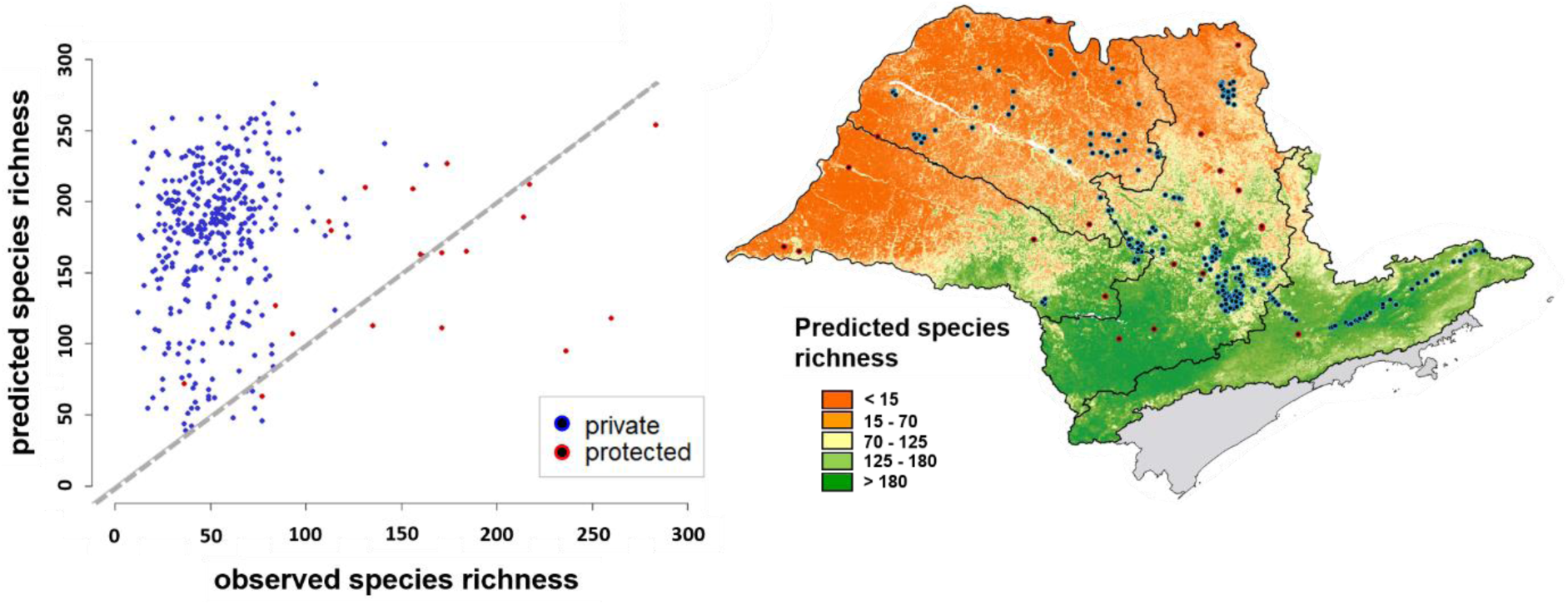
**(a)** Scatterplot relating the predicted and observed species richness at the site level, considering both protection types (strictly Protected Areas and Private lands). Dots below the line represent forest fragments with higher observed richness than predicted. **(b)** Predicted species richness map, calculated by summing the raw suitabilities of occurrence for each of the 663 woody species of this study regions.

Partition of total beta diversity (**β**_**SOR**_) indicated a consistent higher contribution of turnover (**β**_**SIM**_) to overall dissimilarity within regions and protection categories (**Table 2**) (**Appendix 5**).

**Table 2:**
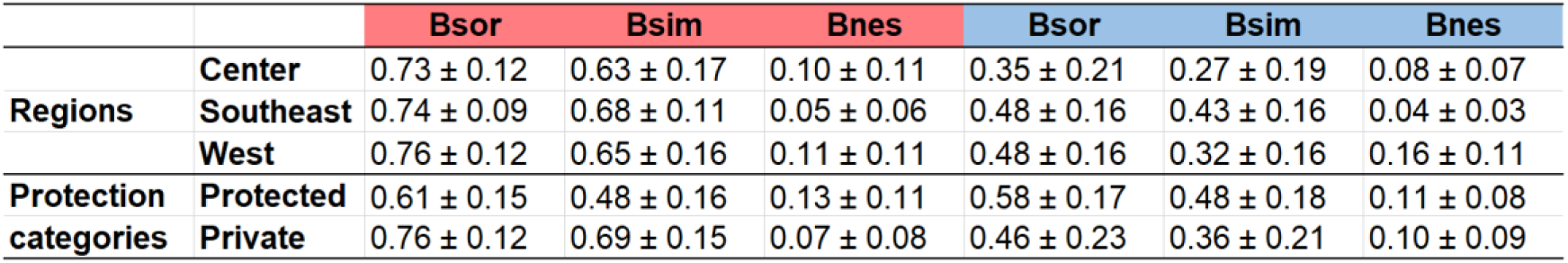
Mean ± standard deviation of total beta diversity (Bsor) and its components turnover (Bsim) and nestedness (Bnes) for distinct regions and categories, considering the observed (red) and the predicted (blue) species composition.

We found that sites with lower predicted than observed **β**_**SOR**_ (ratio < 1) also had higher predicted than observed species richness (ratio > 1) (**Figure 5**), which indicates a correlation between local species loss and heterogenization.

**Figure 5:**
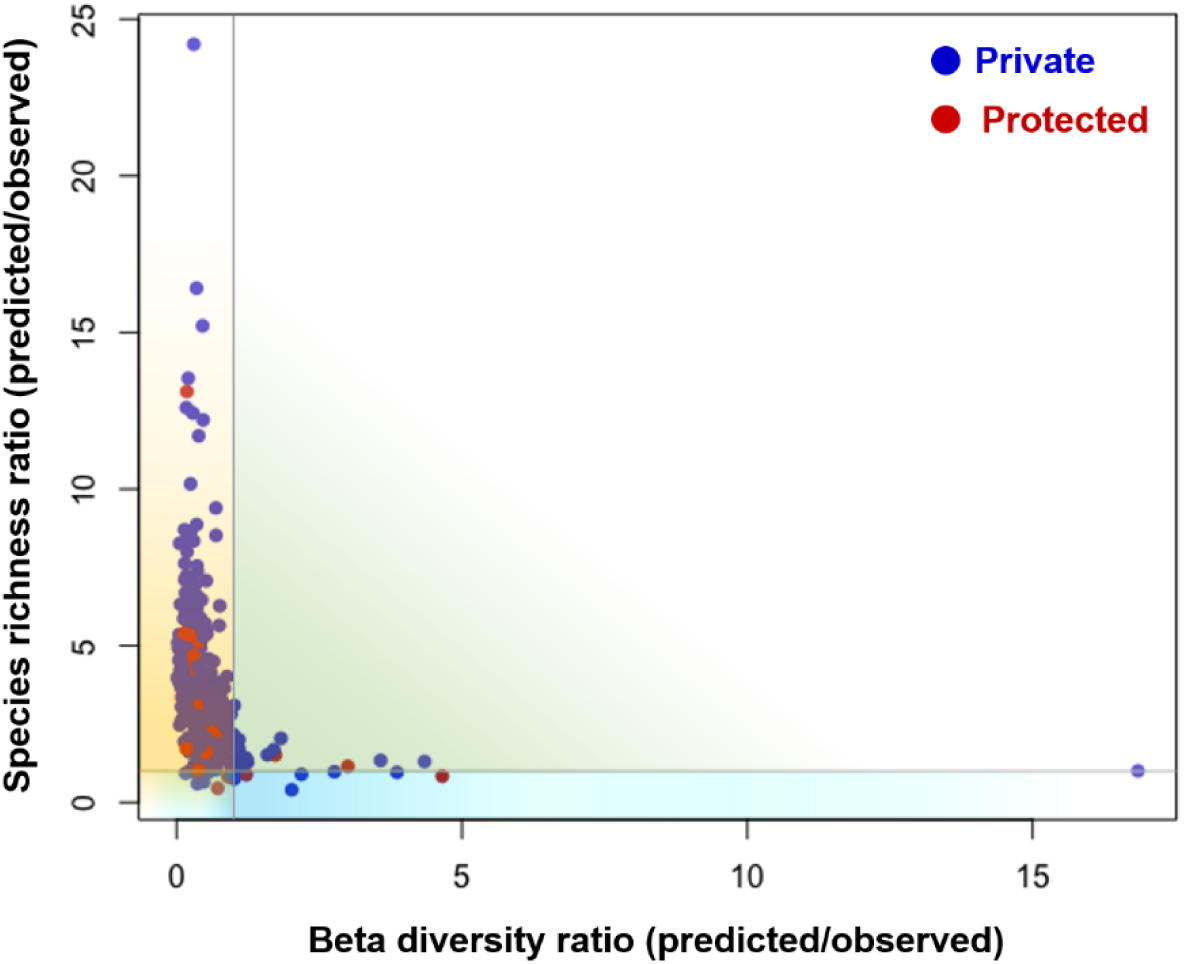
Scatterplot relating the predicted/observed species richness ratio (vertical axis) and total beta diversity (**β**_**SOR**_) ratio (horizontal axis). Sites (dots) above vertical value=1 and below horizontal value=1 (yellow quadrant) experienced local species loss and heterogenization (higher observed **β**_**SOR**_) in comparison to predictions. Green quadrant represent sites that experienced local species loss and homogenization (lower observed **β**_**SOR**_).

The mean observed distance to centroid (**β**_**d**_) was significantly higher than the predicted **β**_**d**_ for all regions and for private lands, with strictly protected areas being the only exception (**Table 3**; **Figure 6**).

**Table 3:** Minimum, mean ± standard deviation and maximum values of distance to centroid (**β**_**d**_) for distinct regions and categories, considering the observed (red) and the predicted (blue) species composition.

| | | Min. | Mean ± SD | Max. | Min. | Mean ± SD | Max. | t-value | P |
| --- | --- | --- | --- | --- | --- | --- | --- | --- | --- |
| Regions | Center | 0.03 | 0.45 ± 0.12 | 0.79 | 0.01 | 0.18 ± 0.15 | 0.72 | 26.18 | <0.0001 |
| | Southeast | 0.08 | 0.47 ± 0.09 | 0.63 | 0.16 | 0.32 ± 0.08 | 0.52 | 9.8 | <0.0001 |
| | West | 0.05 | 0.46 ± 0.10 | 0.65 | 0.02 | 0.22 ± 0.13 | 0.69 | 12.28 | <0.0001 |
| Protection | Protected | 0.16 | 0.37 ± 0.16 | 0.64 | 0.13 | 0.44 ± 0.19 | 0.75 | 0.29 | 0.75 |
| categories | Private | 0.02 | 0.39 ± 0.15 | 0.78 | 0.01 | 0.34 ± 0.20 | 0.73 | 31.79 | <0.0001 |

**Figure 6:**
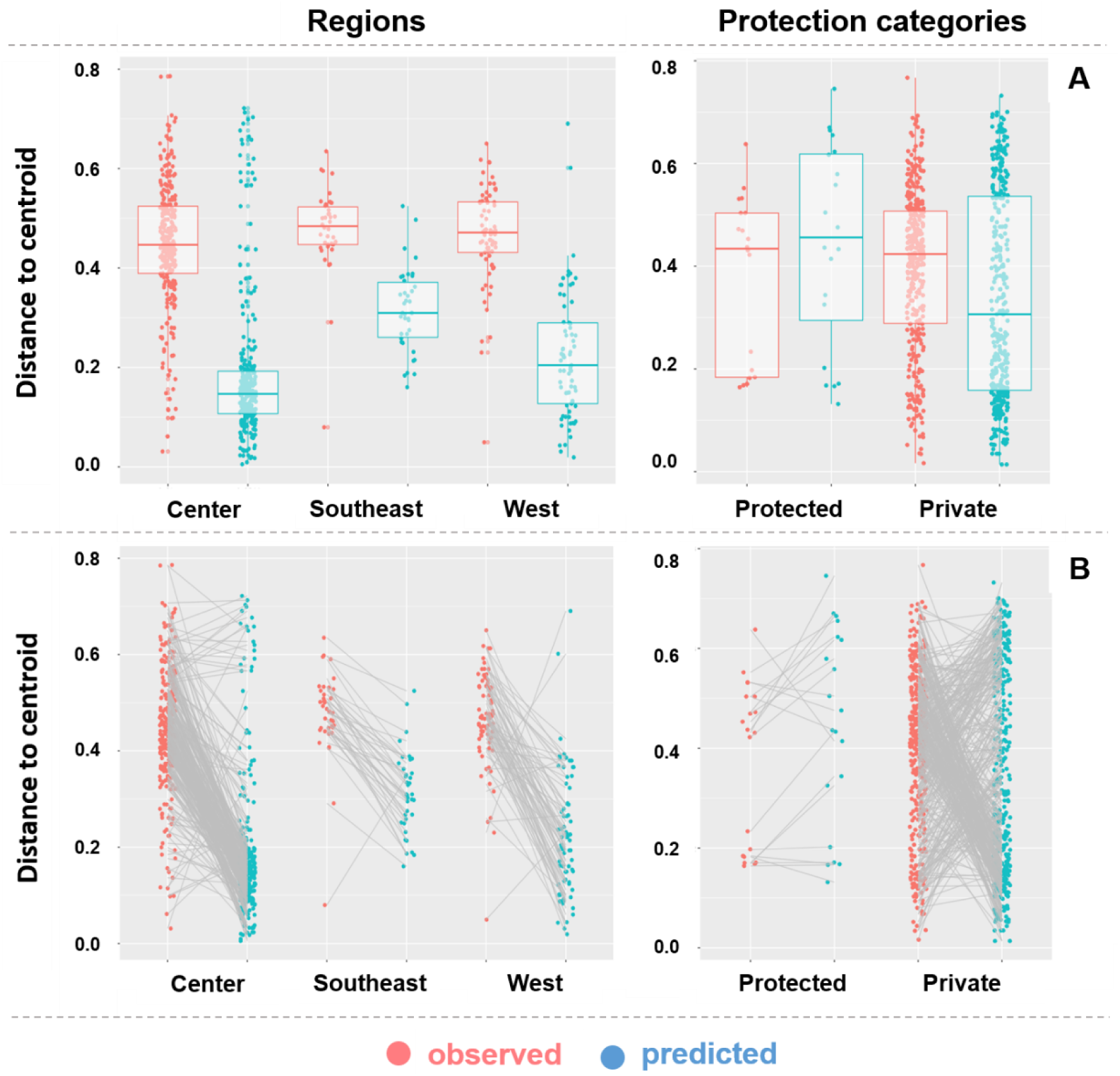
**(a)** Boxplots representing individual distances to centroid (**βd** values) considering the observed (red) and predicted (blue) datasets of different regions and protection categories. Each point represents the distance from one site (forest fragment) to the group centroid in in a βsim-based ordination space. **(b)** Grey lines connect pairs of observed and predicted **βd** values.

Individual **β**_**d**_ values varied within regions and protection categories from almost zero to almost 0.8, especially in the central region (**Figure 6a**). Despite this variation, most observed-predicted pairs of sites showed an increase in **β**_**d**_ from predicted to observed values (**Figure 6b**), indicating a clear trend for biotic heterogenization in the different regions. After splitting the data into protection categories (all regions pooled), we found that the higher beta-diversity in the observed data was due to an increase in **β**_**d**_ values in private lands, as there was no difference between observed and predicted values in strictly protected areas. These results were confirmed by the paired t-test (**Table 3**).

## DISCUSSION

We detected a general loss on observed local diversity in relation to our modeled predictions and this was particularly true among forest fragments in private lands, where we registered consistent reductions in species richness. The higher β diversity registered for the observed dataset imply an overall biotic heterogenization. However, our study also exposes the complexity of this process, with evidence indicating that both homogenization (positive observed-predicted pairwise slopes) and heterogenization (negative observed-predicted pairwise slopes) are taking place in these hyper-fragmented landscapes. Even though distinct regions of São Paulo state were gradually occupied and converted along time – from the eastern coast towards western countryside – all of them presented the same pattern: observed and predicted beta diversity were significantly different, with lower mean values for the latter. This is probably because of the results found among private lands, which represent almost 95% of our samples and drove the pattern registered for all state regions (*i.e.*, heterogenization). As an exception, strictly Protected Areas had lower local species’ loss with no significant differences between observed and predicted β diversity, suggesting they may be fulfilling, to some extent, their protection purpose.

Biodiversity changes in tropical ecosystems are extremely complex to evaluate and understand, as they are scale and context dependent, differ among taxonomic groups and ecosystems, and often respond differently to similar environmental changes (Vellend et al. 2013; Dornelas et al. 2014; Newbold et al. 2015; McGill et al. 2015; Boesing et al. 2018; Catano et al. 2017; Magurran et al. 2018). For instance, recent studies found no evidence for systematic loss in local diversity (Vellend et al. 2013; Dornelas et al. 2014), while several others indicate this is not true for the tropics, where a variety of taxa experienced steep local species decreases in human modified landscapes (Haddad et al. 2015; Mendenhall et al. 2016; Beca et al. 2017; Ceballos et al. 2017; Farah et al. 2017; Galetti et al. 2017; Barlow et al. 2018; Bovendorp et al. 2018). Whereas the unprecedented level of forest degradation, fragmentation and intensive land use have an undeniable contribution to immediate and long-term local diversity loss (Haddad et al. 2015; Barlow et al. 2018), less is known regarding how these disturbances modify and drive the community composition along time and space, regardless of the recent growing interest and evidence on these compositional shifts (Dornelas et al. 2014; Haddad et al. 2015; McGill et al. 2015; Collins et al. 2017; Olden et al. 2018).

The comparison between observed and predicted species composition revealed the idiosyncratic responses of β-diversity (i.e. homogenization and heterogenization) in agricultural landscapes, which can be explained by several mechanisms suggested by the literature. Studies that observed biotic homogenization associate β diversity reduction to niche-selection processes, suggesting ecological filtering overrides environmental heterogeneity (Vellend et al. 2007; Lôbo et al. 2011; Marcelo Tabarelli et al. 2012; Arroyo-Rodríguez et al. 2013; Morante-Filho, Arroyo-Rodríguez, et al. 2015; Püttker et al. 2015; Zwiener et al. 2017). This statement assumes non-random local species extinctions occur because habitat fragmentation affects species differently, according to traits such as rarity, life span, dispersal, and reproductive mode (Haddad et al. 2015), supporting the proliferation of widespread, short-lived and small-seeded species (*e.g.* pioneer species, generalists or “winners” as defined by Tabarelli et al. 2012a) in detriment of rare and shade-tolerant species (*e.g.* specialists or “losers”) (Lôbo et al. 2011; Marcelo Tabarelli et al. 2012; Arroyo-Rodríguez et al. 2013; Morante-Filho, Arroyo-Rodríguez, et al. 2015; Zwiener et al. 2017). Additionally, non-random plant extinctions may also be related to selective logging, which overharvest valuable hardwood species, and to the disappearance of large and medium frugivores through overhunting and habitat loss, with cascading effects over plant-frugivore interactions, species persistence, ecosystem services and functioning in human-modified landscapes (Bello et al. 2015; Bovendorp et al. 2018). All of these mechanistic explanations may be related to the homogenization registered in our study region, where forest fragments are usually small and therefore exposed to edge effects, with depleted plant-animal interactions – especially large-sized species (Beca et al. 2017, Emer et al. 2018), and subject to recurrent fire and other disturbances (Farah et al. 2017).

Overall, however, our results showed that biotic heterogenization is the predominant process in our study region, accordingly to studies that found compositional shifts heading towards divergent communities (Smart et al. 2006; Dornelas et al. 2014; Solar et al. 2015; Sfair et al. 2016; Collins et al. 2017). In fact, a meta-analysis carried by Catano *et al.* (2017) found 21 cases of heterogenization among 22 studies evaluating herbaceous plants in disturbed/undisturbed grasslands and savannas. Assuming the long history of forest degradation and fragmentation in our study region as the main driver acting upon forest fragments, we refer to some mechanisms that may explain why the studied communities were more heterogeneous than compared to our modeled predictions. First, there is a combination of long-term disturbances that impose constant selection pressures (*e.g.*, edge effects) with occasional and contingent perturbations (*e.g.*, fires, windstorms etc.), resulting in unique disturbance histories and shifts in the physical environment (*e.g.* microhabitat conditions), that most likely enhance pre-disturbance compositional differences (Haddad et al. 2015; Catano et al. 2017). Second, forest fragments in private lands are more susceptible to disturbances due to the lack of formal and effective protection, proven by their altogether smaller patch sizes (Ribeiro et al. 2009). They also represent a greater variety of conditions that range from mature forests experiencing post-fragmentation changes to regenerating secondary forests (Laurance et al. 2014; Malhi et al. 2014; Farah et al. 2017). In common, they share reduced local species richness, which together with a large regional species pool (*i.e.*, γ diversity) may create a sampling effect; *i.e.*, a higher probability of more distinct composition between sites when a small portion of the species pool (*i.e.*, low α diversity) is expected to occur in any random community, inflating β diversity (Karp et al. 2012; Newbold et al. 2015). The third explanation is particularly relevant in the studied hyper-fragmented landscapes, where dispersal limitation due to patch isolation might play a dominant role in making those communities such heterogeneous. This is supported by the strong positive relation between local species richness and seed arrival in plant communities (Myers & Harms 2009) and because dispersal limitation play a stronger role in determining community assembly in tropical forests (Myers et al. 2013). More specifically, Catano’s *et al*. (2017) findings on how disturbance and dispersal interact and alter community composition support that increased β-diversity in disturbed landscapes occurs when dispersal is limited, challenging the hypothesis that disturbances always homogenizes communities compositions through deterministic environmental filtering, that is, selecting those species best able to survive within HMLs (Vellend et al. 2007; Lôbo et al. 2011; Arroyo-Rodríguez et al. 2013; Püttker et al. 2015). Finally, other plausible mechanisms acting upon these forest fragments may be related to the reduction in the number of individuals and thus in community size, turning them more susceptible to ecological drift and other stochastic forces (Orrock & Watling 2010), and to competitive release arising from the removal of dominant species (Catano et al. 2017).

Given that, by definition, proper evaluation of biotic homogenization or heterogenization processes depend on quantifying changes in β diversity through space and time (McKinney & Lockwood 1999; Olden & Rooney 2006; Olden et al. 2018), the use of Species Distribution Model proved valuable for predicting community composition in the absence of habitat loss and fragmentation, serving as a temporal surrogate in our study. However, we must acknowledge that our modeling approach imposed some restrictions, notably the non-inclusion of rare or poorly-sampled species and biotic interactions. That said, we do not expect that the overall trend registered here – biotic heterogenization – would be affected by the absence of rare species because their inclusion would most likely increase the differences among communities, while biotic interactions were addressed by choosing a method to adjust or at least reduce an overprediction bias related to the lack of biotic interactions (Calabrese et al. 2014; Gavish et al. 2017; Guisan & Rahbek 2011; D’Amen et al. 2017). Another caveat is that the reduced local species richness among private lands may be related, to some extent, to sampling effort. Since the floristic assessments that compose most of the dataset used here aimed to quickly characterize the regional flora for restoration purposes (Rodrigues et al. 2011), we applied preliminary analysis of incidence-based estimated richness (e.g., Chao 2, Jackknife 1 and Jackknife 2 (Magurran 2013)) that indicated satisfactory sampling effort for both private lands and strictly protected areas. Furthermore, we chose β diversity metrics that focused on compositional changes to alleviate the contribution of α diversity and eventual uneven sampling effort (Koleff et al. 2003). With those considerations, we are confident that our results are consistent and would not be much different from what we have shown here.

Our study highlights the complexity and idiosyncrasies of community compositional shifts in hyper-fragmented landscapes, where both homogenization and heterogenization processes were detected, with the latter prevailing as an overall trend, especially in non-protected private lands. From an applied perspective, the implication of biotic homogenization or heterogenization alone is not sufficient to underpin conservation strategies, as its interpretation is not straightforward – human disturbances can cause β diversity to increase, decrease or remain unchanged (Socolar et al. 2016; Olden et al. 2018). However, the heterogenization process in our study is coupled with a scenario where (i) the originally diverse vegetation is now extremely reduced and fragmented, with small and isolated patches distributed within an intensive agricultural matrix; (ii) forest fragments, especially in private lands, represent unique disturbance histories that result in varying quality habitats, and often in reduced local diversity derived from a large regional species pool (γ diversity); (iii) community composition accumulate great variation among patches (high β diversity), predominantly from turnover (*i.e.* replacement of species). Bringing these facts together and recognizing that conservation of biodiversity extends far beyond the boundaries of strictly Protected Areas, we advocate that all forest fragments are valuable for conservation in HMLs, with particular relevance for private lands, which represent the most exposed and neglected share of what is left (Gardner et al. 2009; Mendenhall et al. 2016; Farah et al. 2017). Based on our results and supported by many other studies, we understand there is enough information to develop an evidence-based approach that should be considered in future management and conservation plans. To foster and sustain biodiversity conservation in HMLs, we thus recommend: (i) effective protection of strictly Protected Areas, which usually represent the largest regional core areas (Joppa et al. 2008) and where compositional shifts apparently are more stable; (ii) active restoration of forest fragments to enhance their alpha diversity, through the management of hyper abundant species (*e.g.*, lianas) (César et al. 2016; Estrada-Villegas & Schnitzer 2018) and reintroduction of lacking groups of species (Garcia et al. 2014; Viani et al. 2015) (iii) active restoration of corridors where the vegetation is degraded and natural regeneration is unlikely, aiming to enhance forest cover and connectivity among forest fragments, allowing species to disperse and persist (Howe 2014; Emer et al. 2018). Finally, considering the growing development of more sustainable agricultural practices (Ferreira et al. 2012, Gonthier et al. 2014) and alternatives for ecological restoration with profitable purposes (Pedro H. S. Brancalion et al. 2012), we encourage the establishment of policies that foster a feasible production model, aligned with the conservation of the remaining biodiversity.

## Author contributions

‘CYV conceived the idea with considerable inputs from RRR, MFS and TS. CYV, MFS and DSBR performed data compilation; MFS and DSBR modeled species distributions; CYV and TS analyzed the data and led the writing of the manuscript; all authors edited the drafts and gave final approval for publication.

## ACKNOWLEDGEMENTS

This study was financed in part by the Coordenação de Aperfeiçoamento de Pessoal de Nível Superior – Brasil (CAPES) – Finance Code 001, by the National Council for Scientific and Technological Development (CNPq grant #140545/2016-6 and #561897/2010-7) and by The São Paulo Research Foundation (FAPESP grant # 2013/50718-5 and #1999/09635-0).

## APPENDICES

**Appendix 1:**
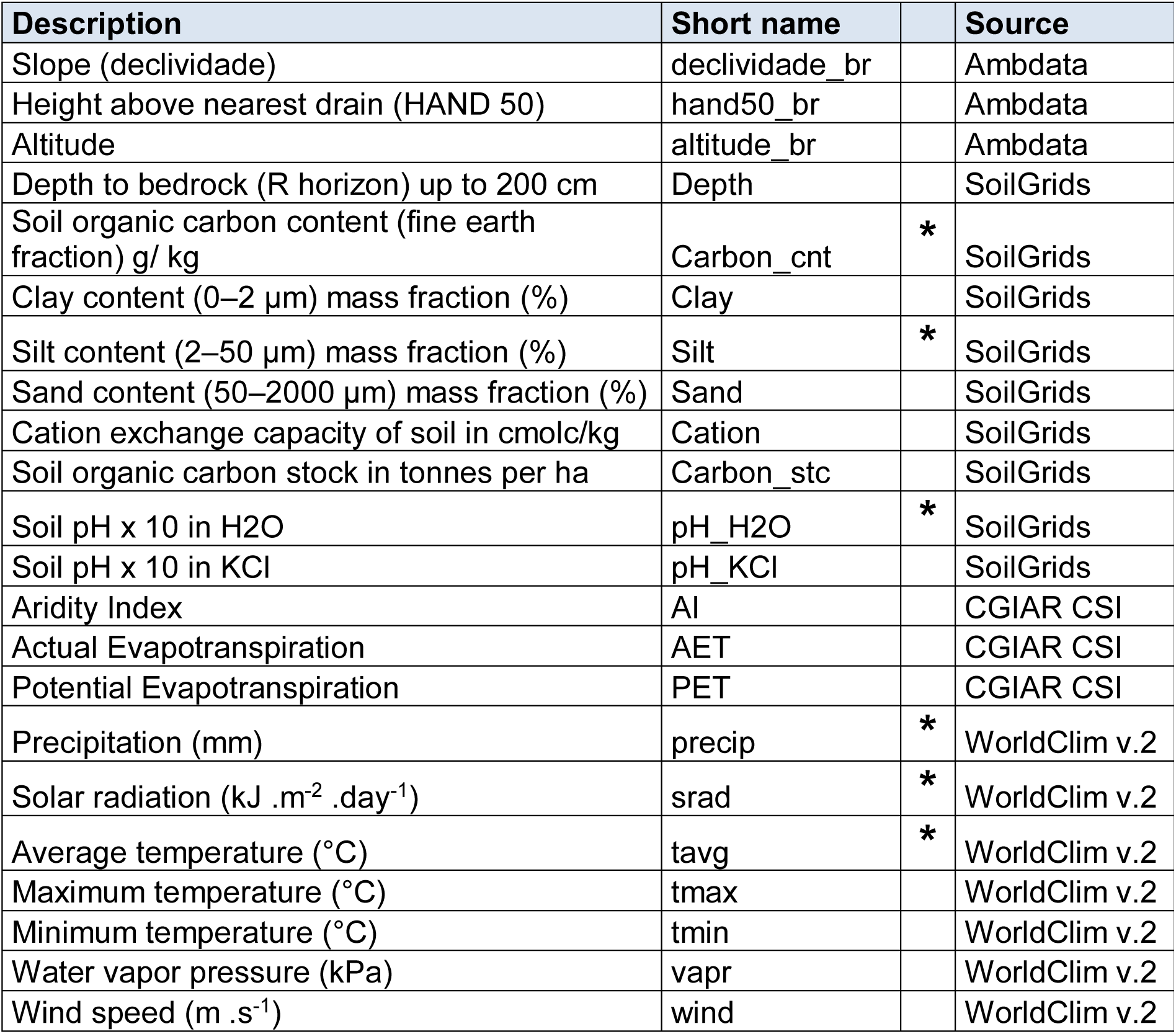
Environmental layers used in the Principal Component Analysis (PCA) and (*) selected for modelling

**Appendix 2:**
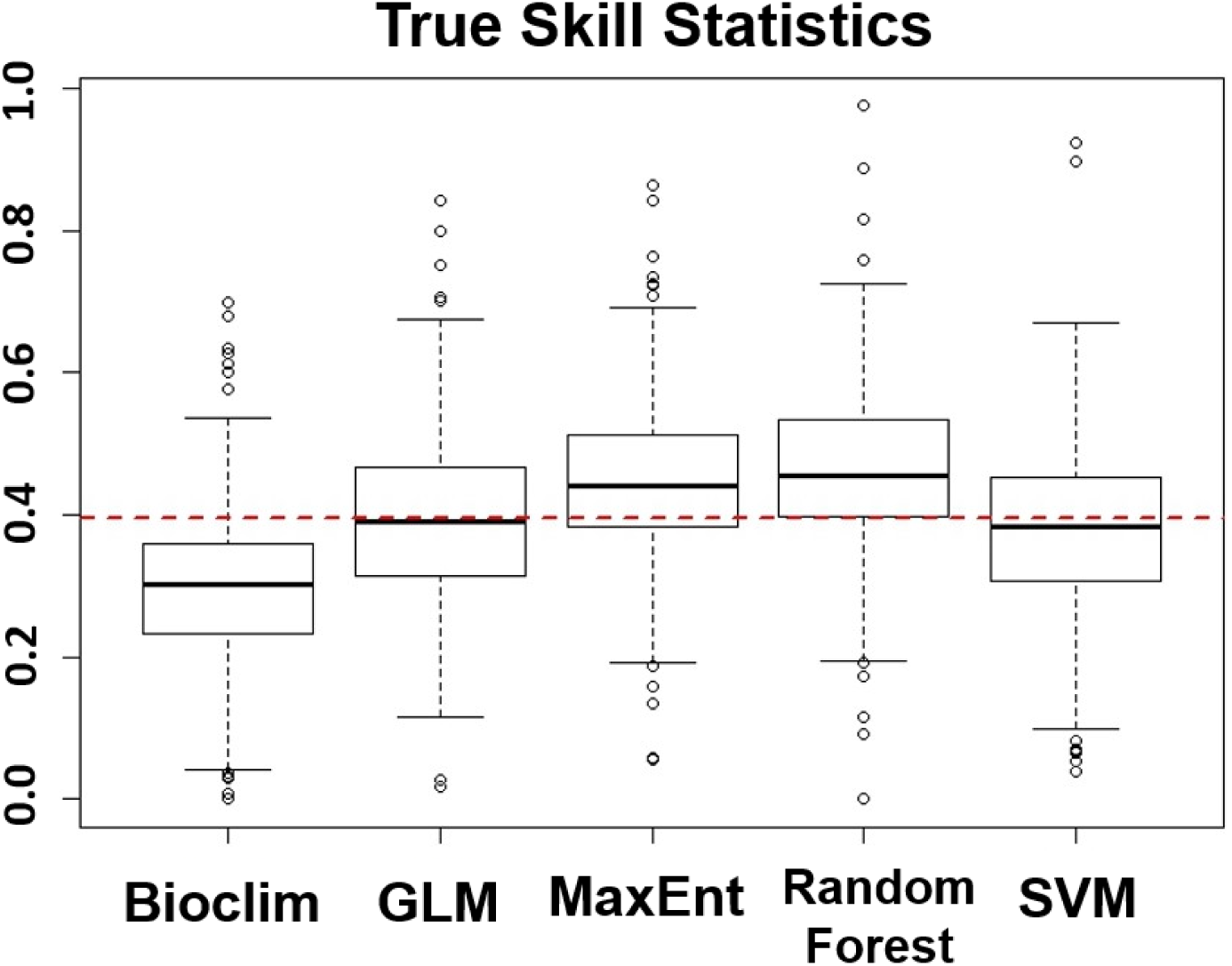
Boxplots (median ± quartiles) of the True Skill Statistics (TSS) comparing partitions of previously tested algorithms used to generate the Species Distribution Models. Dashed red line indicates the selection criteria (TSS=0.4) applied to choose algortihms that composed the final ensemble model (MaxEnt and Random Forest).

**Appendix 3:**
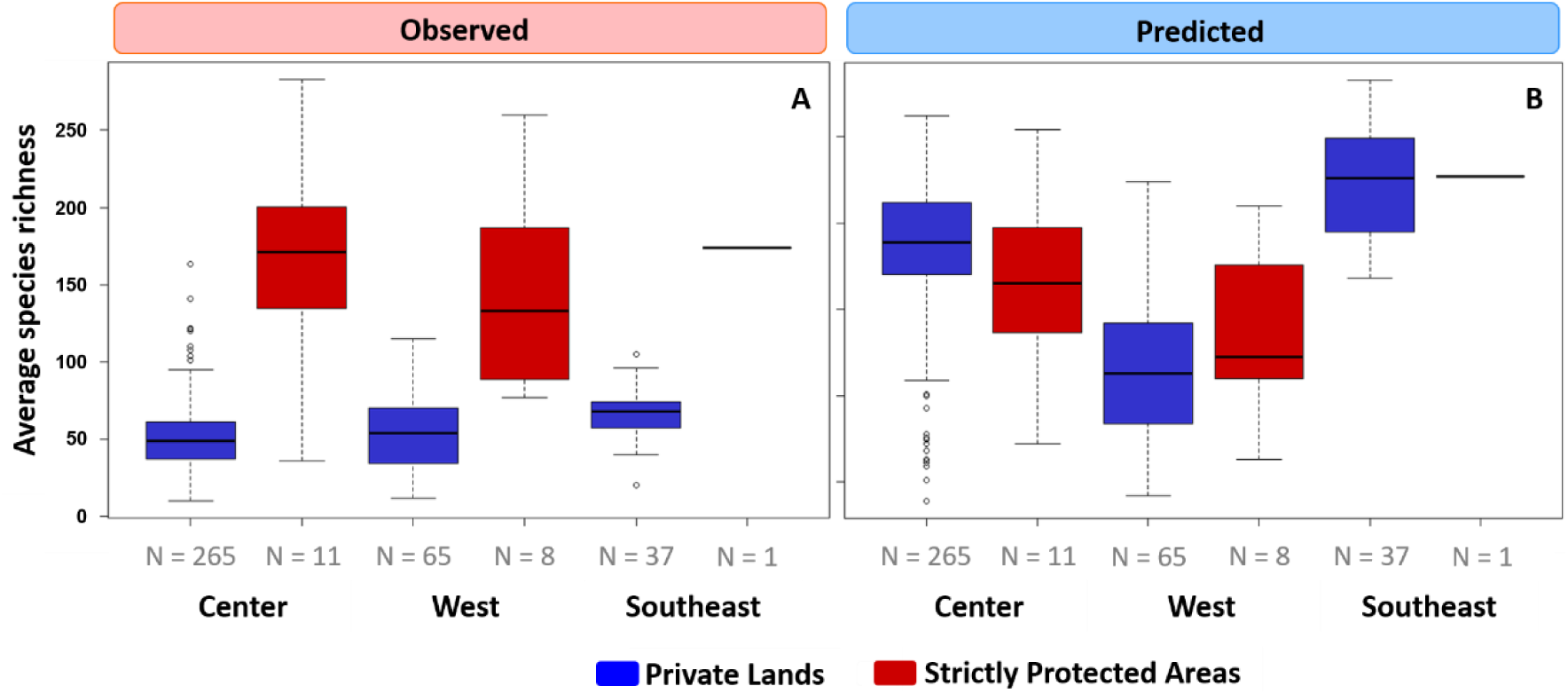
Boxplots (median ± quartiles) comparing protection categories and regions of São Paulo state considering (a) the observed and (b) the predicted species richness per forest fragment. N = sampled forest fragments.

**Appendix 4:**
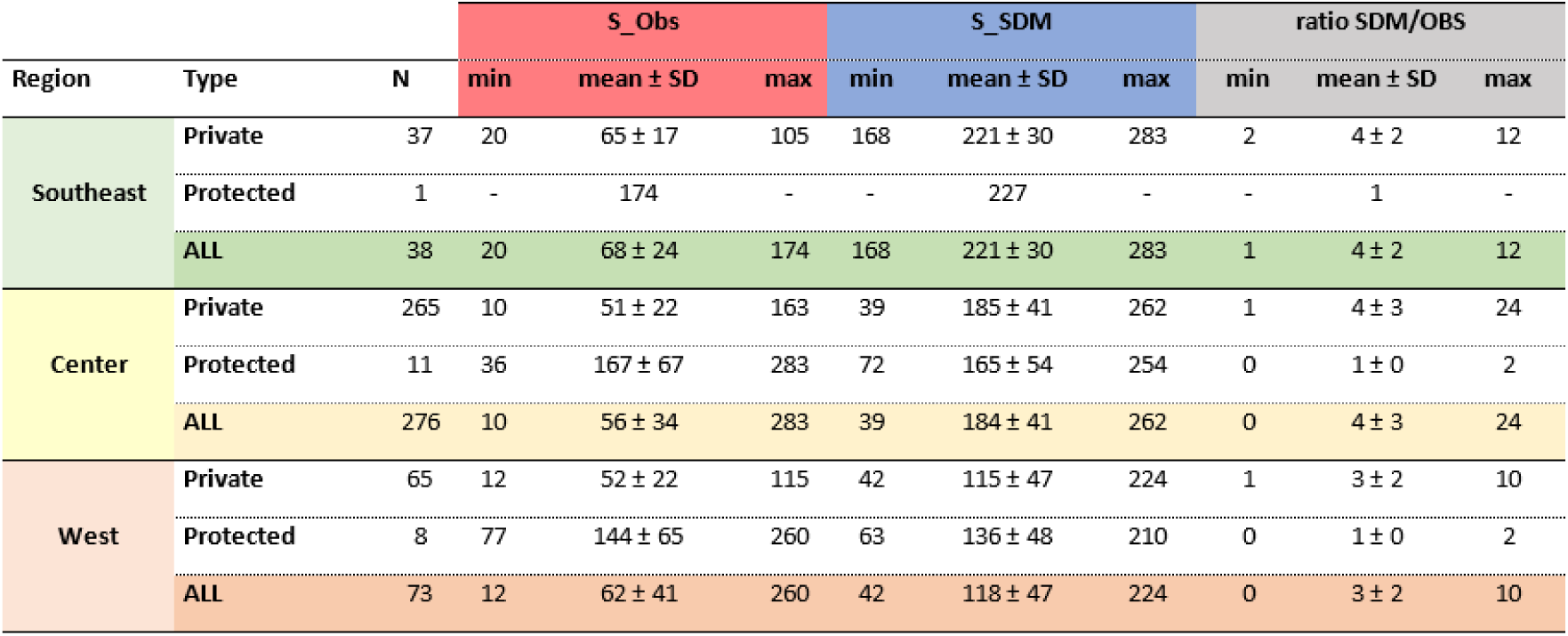
Minimum, mean ± standard deviation and maximum values for species richness for distinct regions and categories, considering the observed (red) and the predicted (blue) species composition, and their ratio (grey). N = sampled forest fragments.

**Appendix 5:**
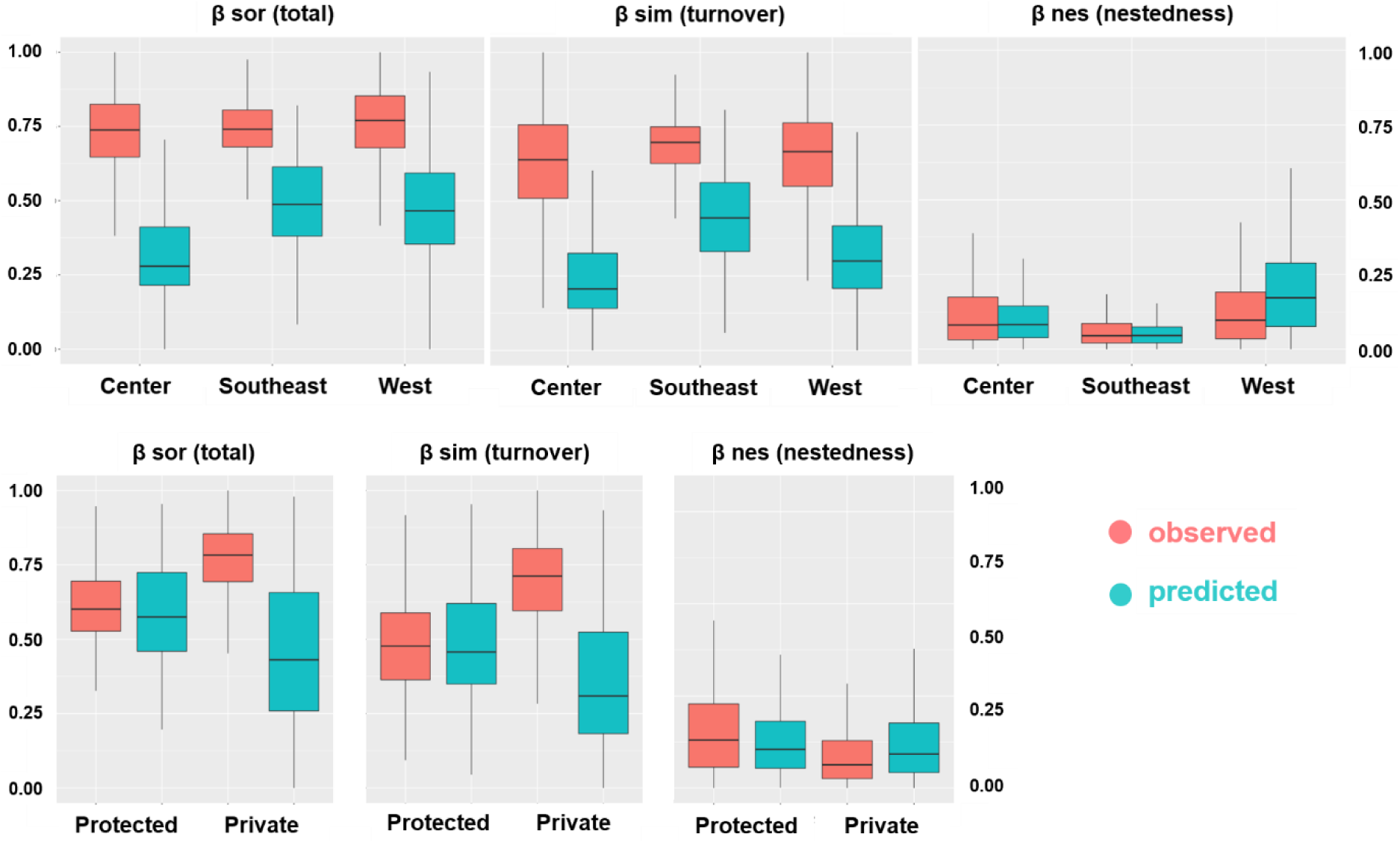
Boxplots (median ± quartiles) of total beta diversity (Bsor) and its components turnover (Bsim) and nestedness (Bnes) for distinct regions and categories, considering the observed (red) and the predicted (blue) species composition.

## REFERENCES

Allouche O, Tsoar A, Kadmon R (2006) Assessing the accuracy of species distribution models: Prevalence, kappa and the true skill statistic (TSS). J Appl Ecol 43:1223–1232. doi: 10.1111/j.1365-2664.2006.01214.x

Andam KS, Ferraro PJ, Pfaff A, et al (2008) Measuring the effectiveness of protected area networks in reducing deforestation. Proc Natl Acad Sci U S A 105:16089–16094. doi: 10.1073/pnas.0800437105

Anderson MJ, Crist TO, Chase JM, et al (2010) Navigating the multiple meanings of β diversity: A roadmap for the practicing ecologist. Ecol Lett 14:19–28. doi: 10.1111/j.1461-0248.2010.01552.x

Anderson MJ, Ellingsen KE, McArdle BH (2006) Multivariate dispersion as a measure of beta diversity. Ecol Lett 9:683–693. doi: 10.1111/j.1461-0248.2006.00926.x

Angelieri CCS, Adams-Hosking C, Ferraz KMPM de B, et al (2016) Using Species Distribution Models to Predict Potential Landscape Restoration Effects on Puma Conservation. PLoS One 11:1–18. doi: 10.1371/journal.pone.0145232

Arroyo-Rodríguez V, Pineda E, Escobar F, Benítez-Malvido J (2008) Value of small patches in the conservation of plant-species diversity in highly fragmented rainforest. Conserv Biol 23:729–39. doi: 10.1111/j.1523-1739.2008.01120.x

Arroyo-Rodríguez V, Rös M, Escobar F, et al (2013) Plant β-diversity in fragmented rain forests: Testing floristic homogenization and differentiation hypotheses. J Ecol 101:1449–1458. doi: 10.1111/1365-2745.12153

Barlow J, França F, Gardner T, et al (2018) The future of hyperdiverse tropical ecosystems. Nature. doi: 10.1038/s41586-018-0301-1

Barlow J, Lennox GD, Ferreira J, et al (2016) Anthropogenic disturbance in tropical forests can double biodiversity loss from deforestation. Nature 1–16. doi: 10.1038/nature18326

Baselga A (2010) Partitioning the turnover and nestedness components of beta diversity. Glob Ecol Biogeogr 19:134–143. doi: 10.1111/j.1466-8238.2009.00490.x

Baselga A, Jiménez-Valverde A, Niccolini G (2007) A multiple-site similarity measure independent of richness. Biol Lett 3:642–645. doi: 10.1098/rsbl.2007.0449

Baselga A, Orme D, Villeger S, et al (2015) betapart: Partitioning beta diversity into turnover and nestedness components. R Packag. Version 1.3 1–26

Beca G, Vancine MH, Carvalho CS, et al (2017) High mammal species turnover in forest patches immersed in biofuel plantations. Biol Conserv. doi: 10.1016/j.biocon.2017.02.033

Bello C, Galetti M, Pizo MA, et al (2015) Defaunation affects carbon storage in tropical forests. Sci Adv 1:1–11. doi: 10.1126/sciadv.1501105

Boesing AL, Nichols E, Metzger JP (2018) Biodiversity extinction thresholds are modulated by matrix type. Ecography (Cop) 1–14. doi: 10.1111/ecog.03365

Bovendorp RS, Brum FT, McCleery RA, et al (2018) Defaunation and fragmentation erode small mammal diversity dimensions in tropical forests. Ecography (Cop). doi: 10.1111/ecog.03504

Brancalion PHS, Viani RAG, Strassburg BBN, Rodrigues RR (2012) Finding the money for tropical forest restoration. Unasylva 63:41–50

Calabrese JM, Certain G, Kraan C, Dormann CF (2014) Stacking species distribution models and adjusting bias by linking them to macroecological models. Glob Ecol Biogeogr 23:99–112. doi: 10.1111/geb.12102

Carranza T, Balmford A, Kapos V, Manica A (2014) Protected area effectiveness in reducing conversion in a rapidly vanishing ecosystem: The Brazilian Cerrado. Conserv Lett 7:216–223. doi: 10.1111/conl.12049

Carvalho GH, Cianciaruso MV, Batalha MA (2010) Plantminer: A web tool for checking and gathering plant species taxonomic information. Environ Model Softw 25:815–816. doi: 10.1016/j.envsoft.2009.11.014

Catano CP, Dickson TL, Myers JA (2017) Dispersal and neutral sampling mediate contingent effects of disturbance on plant beta-diversity: a meta-analysis. Ecol Lett 20:347–356. doi: 10.1111/ele.12733

Ceballos G, Ehrlich PR, Dirzo R (2017) Biological annihilation via the ongoing sixth mass extinction signaled by vertebrate population losses and declines. Proc Natl Acad Sci 201704949. doi: 10.1073/pnas.1704949114

César RG, Holl KD, Girão VJ, et al (2016) Evaluating climber cutting as a strategy to restore degraded tropical forests. Biol Conserv 201:309–313. doi: 10.1016/j.biocon.2016.07.031

Chao A, Chiu CH, Hsieh TC, Inouye BD (2012) Proposing a resolution to debates on diversity partitioning. Ecology 93:2037–2051. doi: 10.1890/11-1817.1

Chazdon RL, Harvey CA, Komar O, et al (2009a) Beyond Reserves?: A Research Agenda for Conserving Biodiversity in Human-modified Tropical Landscapes. Biotropica 41:142–153

Chazdon RL, Peres C a, Dent D, et al (2009b) The potential for species conservation in tropical secondary forests. Conserv Biol 23:1406–17. doi: 10.1111/j.1523-1739.2009.01338.x

Coetzee BWT, Gaston KJ, Chown SL (2014) Local scale comparisons of biodiversity as a test for global protected area ecological performance: A meta-analysis. PLoS One 9:. doi: 10.1371/journal.pone.0105824

Collins CD, Banks-Leite C, Brudvig LA, et al (2017) Fragmentation affects plant community composition over time. Ecography (Cop) 40:119–130. doi: 10.1111/ecog.02607

D’Amen M, Pradervand J-N, Guisan A (2015) Predicting richness and composition in mountain insect communities at high resolution?: a new test of the SESAM framework. Glob Ecol Biogeogr 24:1443–1453. doi: 10.1111/geb.12357

D’Amen MD, Rahbek C, Zimmermann NE, Guisan A (2017) Spatial predictions at the community level?: from current approaches to future frameworks. Biol Rev 92:169–187. doi: 10.1111/BRV.12222

De Marco Junior P, Siqueira MF (2009) Como determinar a distribuição potencial de espécies sob uma abordagem conservacionista? Megadiversidade 5:65–76

Dent DH, Joseph Wright S (2009) The future of tropical species in secondary forests: A quantitative review. Biol Conserv 142:2833–2843. doi: 10.1016/j.biocon.2009.05.035

Diniz-filho AF, Bini LM, Rangel TF, et al (2009) Partitioning and mapping uncertainties in ensembles of forecasts of species turnover under climate change. Ecography (Cop) 32:897–906. doi: 10.1111/j.1600-0587.2009.06196.x

Dornelas M, Gotelli NJ, McGill B, et al (2014) Assemblage Time Series Reveal Biodiversity Change but Not Systematic Loss. Science (80-) 344:296–299. doi: 10.1126/science.1248484

Durigan G, Ratter JA (2006) Successional changes in cerrado and cerrado/forest ecotonal vegetation in western São Paulo State, Brazil, 1962-2000. Edinburgh J Bot 63:119–130. doi: 10.1017/S0960428606000357

Durigan G, Siqueira MF, Franco GADC, Ratter JA (2006) Seleção De Fragmentos Prioritários Para a Criação De Unidades De Conservação Do Cerrado No Estado De São Paulo. Rev do Inst Florest 18:23–37. doi: 10.1007/s13398-014-0173-7.2

Elith J, Graham C, Anderson R, et al (2006) Novel methods improve prediction of species’ distributions from occurrence data. Ecography (Cop) 29:129–151. doi: 10.1111/j.2006.0906-7590.04596.x

Emer C, Galetti M, Pizo MA, et al (2018) Seed-dispersal interactions in fragmented landscapes – a metanetwork approach. Ecol Lett 21:484–493. doi: 10.1111/ele.12909

Estrada-Villegas S, Schnitzer SA (2018) A comprehensive synthesis of liana removal experiments in tropical forests. Biotropica 0:1–11. doi: 10.1111/btp.12571

Farah FT, Muylaert R de L, Ribeiro MC, et al (2017) Integrating plant richness in forest patches can rescue overall biodiversity in human-modified landscapes. For Ecol Manage 397:78–88

Flora_do_Brasil_2020 Jardim Botânico do Rio de Janeiro. Available on http://floradobrasil.jbrj.gov.br/

Galetti M, Brocardo CR, Begotti RA, et al (2017) Defaunation and biomass collapse of mammals in the largest Atlantic forest remnant. Anim Conserv 20:270–281. doi: 10.1111/acv.12311

Galetti M, Giacomini HC, Bueno RS, et al (2009) Priority areas for the conservation of Atlantic forest large mammals. Biol Conserv 142:1229–1241. doi: 10.1016/j.biocon.2009.01.023

Garcia LC, Hobbs RJ, Mäes dos Santos F a., Rodrigues RR (2014) Flower and Fruit Availability along a Forest Restoration Gradient. Biotropica 46:114–123. doi: 10.1111/btp.12080

Gardner T a., Barlow J, Chazdon R, et al (2009) Prospects for tropical forest biodiversity in a human-modified world. Ecol Lett 12:561–582. doi: 10.1111/j.1461-0248.2009.01294.x

Gavish Y, Marsh CJ, Kuemmerlen M, et al (2017) Accounting for biotic interactions through alpha-diversity constraints in stacked species distribution models. Methods Ecol Evol. doi: 10.1111/2041-210X.12731

Guisan A, Rahbek C (2011) SESAM – a new framework integrating macroecological and species distribution models for predicting spatio-temporal patterns of species assemblages. J Biogeogr 38:1433–1444. doi: 10.1111/j.1365-2699.2011.02550.x

Guisan A, Tingley R, Baumgartner JB, et al (2013) Predicting species distributions for conservation decisions. Ecol Lett 16:1424–1435. doi: 10.1111/ele.12189

Haddad NM, Brudvig LA, Clobert J, et al (2015) Habitat fragmentation and its lasting impact on Earth’s ecosystems. Science (80-) 1–9

Hansen MC, Potapov P V, Moore R, et al (2013) High-resolution global maps of 21stcentury forest cover change. Science 342:850–3. doi: 10.1126/science.1244693

Howe HF (2014) Diversity Storage: Implications for tropical conservation and restoration. Glob Ecol Conserv 2:349–358. doi: 10.1016/j.gecco.2014.10.004

Joppa LN, Loarie SR, Pimm SL (2008) On the protection of “protected areas.” Proc Natl Acad Sci 105:6673–6678

Jost L (2007) Partitioning diversity into independent alpha and beta components. Ecology 88:2427–2439. doi: 10.1890/06-1736.1

Jost L, Chao A, Chazdon RL (2010) Compositional similarity and B (beta) diversity. In: Biological diversity: frontiers in measurement and assessment. p 368

Karp DS, Rominger AJ, Zook J, et al (2012) Intensive agriculture erodes B-diversity at large scales. Ecol Lett 15:963–970. doi: 10.1111/j.1461-0248.2012.01815.x

Koleff P, Gaston KJ, Lennon JJ (2003) Measuring beta diversity for presence – absence data. J Anim Ecol 72:367–382. doi: 10.1046/j.1365-2656.2003.00710.x

Kurtz BC, Valentin JL, Scarano FR (2015) Are the Neotropical Swamp Forests a Distinguishable Forest Type? Patterns From Southeast and Southern Brazil. Edinburgh J Bot 72:191–208. doi: 10.1017/S096042861400033X

Laurance WF (2009) Conserving the hottest of the hotspots. Biol Conserv 142:1137. doi: 10.1016/j.biocon.2008.10.011

Laurance WF, Sayer J, Cassman KG (2014) Agricultural expansion and its impacts on tropical nature. Trends Ecol Evol 29:107–116. doi: 10.1016/j.tree.2013.12.001

Legendre P, De Cáceres M (2013) Beta diversity as the variance of community data: Dissimilarity coefficients and partitioning. Ecol Lett 16:951–963. doi: 10.1111/ele.12141

Lima RAF de, Mori DP, Pitta G, et al (2015a) How much do we know about the endangered Atlantic Forest? Reviewing nearly 70 years of information on tree community surveys. Biodivers Conserv. doi: 10.1007/s10531-015-0953-1

Lima PB, Lima LF, Santos B a., et al (2015b) Altered herb assemblages in fragments of the Brazilian Atlantic forest. Biol Conserv 191:588–595. doi: 10.1016/j.biocon.2015.08.014

Lôbo D, Leão T, Melo FPL, et al (2011) Forest fragmentation drives Atlantic forest of northeastern Brazil to biotic homogenization. Divers Distrib 17:287–296. doi: 10.1111/j.1472-4642.2010.00739.x

Loyola R (2014) Brazil cannot risk its environmental leadership. Divers Distrib 20:1365–1367. doi: 10.1111/ddi.12252

Magurran AE (2013) Measuring biological diversity

Magurran AE, Deacon AE, Moyes F, et al (2018) Divergent biodiversity change within ecosystems. Proc Natl Acad Sci 201712594. doi: 10.1073/pnas.1712594115

Malhi Y, Gardner T a., Goldsmith GR, et al (2014) Tropical Forests in the Anthropocene. Annu Rev Environ Resour 39:125–159. doi: 10.1146/annurev-environ-030713-155141

Martinelli LA, Joly CA, Nobre CA, Sparovek G (2010) A falsa dicotomia entre a preservação da vegetação natural e a produção agropecuária. Biota Neotrop 10:323–330. doi: 10.1590/S1676-06032010000400036

McGill BJ, Dornelas M, Gotelli NJ, Magurran AE (2015) Fifteen forms of biodiversity trend in the anthropocene. Trends Ecol Evol 30:104–113. doi: 10.1016/j.tree.2014.11.006

McKinney ML, Lockwood JL (1999) Biotic homogenization: A few winners replacing many losers in the next mass extinction. Trends Ecol Evol 14:450–453. doi: 10.1016/S0169-5347(99)01679-1

Melo FPL, Arroyo-Rodríguez V, Fahrig L, et al (2013) On the hope for biodiversityfriendly tropical landscapes. Trends Ecol Evol 28:462–8. doi: 10.1016/j.tree.2013.01.001

Mendenhall CD, Shields-Estrada A, Krishnaswami AJ, Daily GC (2016) Quantifying and sustaining biodiversity in tropical agricultural landscapes. Proc Natl Acad Sci 113:14544–14551. doi: 10.1073/pnas.1604981113

Metzger JP (2009) Conservation issues in the Brazilian Atlantic forest. Biol Conserv 142:1138–1140. doi: 10.1016/j.biocon.2008.10.012

Morante-Filho JC, Arroyo-Rodríguez V, Faria D (2015a) Patterns and predictors of beta diversity in the fragmented Brazilian Atlantic forest: A multiscale analysis of forest specialist and generalist birds. J Anim Ecol 85:240–250. doi: 10.1111/1365-2656.12448

Myers J a., Harms KE (2009) Seed arrival, ecological filters, and plant species richness: A meta-analysis. Ecol Lett 12:1250–1260. doi: 10.1111/j.1461-0248.2009.01373.x

Myers JA, Chase JM, Jiménez I, et al (2013) Beta-diversity in temperate and tropical forests reflects dissimilar mechanisms of community assembly. Ecol Lett 16:151–157. doi: 10.1111/ele.12021

Myers N, Mittermeier RA, Mittermeier CG, et al (2000) Biodiversity hotspots for conservation priorities. Nature 403:853–858. doi: 10.1038/35002501

Newbold T, Hudson LN, Hill SLL, et al (2015) Global effects of land use on local terrestrial biodiversity. Nature. doi: 10.1038/nature14324

Olden JD, Comte L, Giam X (2018) The Homogocene: a research prospectus for the study of biotic homogenisation. NeoBiota 37:23–36. doi: 10.3897/neobiota.37.22552

Olden JD, Rooney TP (2006) On defining and quantifying biotic homogenization. Glob Ecol Biogeogr 15:113–120. doi: 10.1111/j.1466-822X.2006.00214.x

Oliveira-Filho A, Fontes M (2000) Patterns of Floristic Differentiation among Atlantic Forests in Southeastern Brazil and the Influence of Climate1. Biotropica 32:793–810. doi: 10.1111/j.1744-7429.2000.tb00619.x

Oliveira-Filho A (2017) NeoTropTree, Flora Arbórea da Região Neotropical: Um Banco de Dados Envolvendo Biogeografia, Diversidade e Conservação (Universidade Federal de Minas Gerais, 2017); http://www.neotroptree.info/

Orrock JL, Watling JI (2010) Local Community size mediates ecological drift and competition in metacommunities. Proc R Soc B Biol Sci 277:2185–2191. doi: 10.1098/rspb.2009.2344

Pardini R, Bueno ADA, Gardner T a, et al (2010) Beyond the fragmentation threshold hypothesis: regime shifts in biodiversity across fragmented landscapes. PLoS One 5:e13666. doi: 10.1371/journal.pone.0013666

Peterson AT, Soberon J (2012) Species Distribution Modeling and Ecological Niche Modeling: Getting the Concepts Right. Nat Conserv 10:102–107. doi: 10.4322/natcon.2012.019

Püttker T, Bueno A de A, Prado PI, Pardini R (2015) Ecological filtering or random extinction? Beta-diversity patterns and the importance of niche-based and neutral processes following habitat loss. Oikos 124:206–215. doi: 10.1111/oik.01018

Ribeiro MC, Metzger JP, Martensen AC, et al (2009) The Brazilian Atlantic Forest: How much is left, and how is the remaining forest distributed? Implications for conservation. Biol Conserv 142:1141–1153. doi: 10.1016/j.biocon.2009.02.021

Rodrigues RR, Gandolfi S, Nave AG, et al (2011) Large-scale ecological restoration of high-diversity tropical forests in SE Brazil. For Ecol Manage 261:1605–1613. doi: 10.1016/j.foreco.2010.07.005

Sánchez-Tapia A, de Siqueira MF, Lima RO, et al (2018) Model-R: A framework for scalable and reproducible ecological niche modeling. Commun Comput Inf Sci 796:218–232. doi: 10.1007/978-3-319-73353-1_15

Santos K, Kinoshita LS, Santos FAM (2007) Tree species composition and similarity in semideciduous forest fragments of southeastern Brazil. Biological Conservation 135(2): 268–277

Setzer J (1966) Atlas climático e ecológico do estado de São Paulo. 61

Sfair JC, Arroyo-Rodriguez V B. A. S, Tabarelli M (2016) Taxonomic and functional divergence of tree assemblages in a fragmented tropical forest. Ecol Appl 2:1816–1826. doi: 10.1890/15-1673.1

Slik JWF, Arroyo-Rodríguez V, Aiba S-I, et al (2015) An estimate of the number of tropical tree species. Proc Natl Acad Sci 112:7472–7477. doi: 10.1073/pnas.1512611112

Smart SM, Thompson K, Marrs RH, et al (2006) Biotic homogenization and changes in species diversity across human-modified ecosystems. Proc Biol Sci 273:2659–2665. doi: 10.1098/rspb.2006.3630

Soares-Filho B, Rajão R, Macedo M, et al (2014) Cracking Brazil’s Forest Code. Science (80-) 344:363–364

Socolar JB, Gilroy JJ, Kunin WE, Edwards DP (2016) How Should Beta-Diversity Inform Biodiversity Conservation? Trends Ecol Evol 31:67–80. doi: 10.1016/j.tree.2015.11.005

Solar RRDC, Barlow J, Ferreira J, et al (2015) How pervasive is biotic homogenization in human-modified tropical forest landscapes? Ecol Lett n/a-n/a. doi: 10.1111/ele.12494

Sparovek G, Berndes G, Barretto AGDOP, Klug ILF (2012) The revision of the brazilian forest act: Increased deforestation or a historic step towards balancing agricultural development and nature conservation? Environ Sci Policy 16:65–72. doi: 10.1016/j.envsci.2011.10.008 *SpeciesLink* (SpeciesLink, accessd 15 March 2017); http://splink.cria.org.br/

Tabarelli M, Aguiar AV, Ribeiro MC, et al (2010) Prospects for biodiversity conservation in the Atlantic Forest: Lessons from aging human-modified landscapes. Biol Conserv 143:2328–2340. doi: 10.1016/j.biocon.2010.02.005

Tabarelli M, Peres C a., Melo FPL (2012a) The “few winners and many losers” paradigm revisited: Emerging prospects for tropical forest biodiversity. Biol Conserv 155:136–140. doi: 10.1016/j.biocon.2012.06.020

Tabarelli M, Santos B a., Arroyo-rodríguez V, Melo F (2012b) Secondary forests as biodiversity repositories in human-modified landscapes: insights from the Neotropics. Bol do Mus do Pará Emílio Goeldi 7:319–328

Tuomisto H (2010) A diversity of beta diversities: Straightening up a concept gone awry. Part 1. Defining beta diversity as a function of alpha and gamma diversity. Ecography (Cop) 33:2–22. doi: 10.1111/j.1600-0587.2009.05880.x

Vellend M, Baeten L, Myers-Smith IH, et al (2013) Global meta-analysis reveals no net change in local-scale plant biodiversity over time. Proc Natl Acad Sci U S A 110:19456–9. doi: 10.1073/pnas.1312779110

Vellend M, Verheyen K, Flinn KM, et al (2007) Homogenization of forest plant communities and weakening of species – environment relationships via agricultural land use. J Ecol 95:565–573. doi: 10.1111/j.1365-2745.2007.01233.x

Viani RAG, Mello FNA, Chi IE, Brancalion PHS (2015) A new focus for ecological restoration: management of degraded forest remnants in fragmented landscapes. GPL news november:5–9

Zwiener VP, Lira-Noriega A, Grady CJ, et al (2017) Climate change as a driver of biotic homogenization of woody plants in the Atlantic Forest. Glob Ecol Biogeogr 1–12. doi: 10.1111/geb.12695

